# Decoding violated sensory expectations from the auditory cortex of anaesthetized mice: Hierarchical recurrent neural network depicts separate ‘danger’ and ‘safety’ units

**DOI:** 10.1101/2022.04.29.490005

**Authors:** Jamie A. O’Reilly, Thanate Angsuwatanakul, Jordan Wehrman

## Abstract

The ability to respond appropriately to sensory information received from the external environment is among the most fundamental capabilities of central nervous systems. In the auditory domain, processes underlying this behaviour are studied by measuring auditory-evoked electrophysiology during sequences of sounds with predetermined regularities. Identifying neural correlates of ensuing auditory novelty responses is supported by research in experimental animals. In the present study, we reanalysed epidural field potential recordings from the auditory cortex of anaesthetised mice during frequency and intensity oddball stimulation. Multivariate pattern analysis (MVPA) and hierarchical recurrent neural network (RNN) modelling were adopted to explore these data with greater resolution than previously considered using conventional methods. Time-wise and generalised temporal decoding MVPA approaches revealed previously underestimated asymmetry between responses to sound-level transitions in the intensity oddball paradigm, in contrast with tone frequency changes. After training, the cross-validated RNN model architecture with four hidden layers produced output waveforms in response to simulated auditory inputs that were strongly correlated with grand-average auditory-evoked potential waveforms (r^2^ > 0.9). Units in hidden layers were classified based on their temporal response properties and characterised using principal component analysis and sample entropy. These demonstrated spontaneous alpha rhythms, sound onset and offset responses, and putative ‘safety’ and ‘danger’ units activated by relatively inconspicuous and salient changes in auditory inputs, respectively. The hypothesised existence of corresponding biological neural sources is naturally derived from this model. If proven, this would have significant implications for prevailing theories of auditory processing.

## 1. Introduction

The survival of humans and animals relies on their ability to detect and respond appropriately to the environment. This is a fundamental behaviour shared by many species. When residing in a dark forest, for example, the mammalian central auditory system automatically processes salient changes in the acoustic environment and redirects the organism’s attention towards any potential sources of danger or reward. These processes can be studied by recording electrophysiological responses to expected and unexpected sounds in a passive auditory oddball sequence. In humans, this is typically performed while recording electroencephalography (EEG), after which averaging responses measured over multiple presentations of the same stimulus condition produces a stereotypical pattern of components in the event-related potential (ERP). These auditory novelty responses are interpreted to reflect surprise or prediction-error signalling, and include mismatch negativity (MMN), P3a and reorienting negativity (RON), which are distinct from obligatory components of the auditory-evoked response that are elicited by all perceptible stimuli, regardless of their context (e.g. P50 or N1).

One of the most widely studied ERP components related to surprise is the MMN (Näätänen et al., 1978). This component occurs between 100 and 200 ms after the onset of a surprising stimulus at fronto-central electrodes (Näätänen et al., 2007) and is calculated from the difference between the responses to expected ‘standard’ and unexpected ‘deviant’ stimuli. The larger the MMN, the greater the level of surprise (De-Wit et al., 2010; Friston and Kiebel, 2009; Garrido et al., 2009; Heilbron and Chait, 2018; Mathys et al., 2011). Supporting this, MMN has been found to be relatively diminished under several conditions in which sensory predictions are thought to be compromised, for example in schizophrenia (Umbricht and Krljes, 2005), Parkinson’s disease (Brønnick et al., 2010) and when subjects are administered ketamine (Rosburg and Kreitschmann-Andermahr, 2016). It has also be observed in unresponsive patients, and there is evidence supporting the effectiveness of both MMN and P3a in evaluating conserved physiological function in unconscious humans (Morlet and Fischer, 2014).

The P3a is a positive response to more surprising stimuli around 250 to 300 ms after the stimulus (Friedman et al., 2001; Polich, 2007). While the MMN is an early index of surprise, the P3a is thought to reflect the attention directed towards the surprising stimulus (Friedman et al., 2001). A loud growling noise in the forest, for example, would likely draw our attention, resulting in a larger P3a. However, if the growling noise is quickly determined to be that of a smaller, non-threatening source, our attention would return to our current task. This reorientation to prior task elicits a much later negative ERP component, the RON. As per the P3a and MMN, this response occurs at fronto-central electrodes, though much later, around 400 to 600 ms post-stimulus (Otten et al., 2000; Schröger and Wolff, 1998). Similar to the MMN, both the P3a and RON are affected by neurophysiological deficits, for example, in schizophrenia patients (Higuchi et al., 2014). Further, it is worth noting that, while related, each of these three components may be separated experimentally (Horváth et al., 2008).

Neural activity associated with these processes is examined in animal models on the basis that their auditory systems are homologous to those in humans. It remains challenging, however, to directly associate signals measured from animals with equivalent components observed in human ERP waveforms, due to anatomical and physiological differences. Ambiguity thereby arises concerning the latency of obligatory and context-dependent components of the auditory response in animals, which may be problematic to dissociate (Parras et al., 2017; Taaseh et al., 2011). Some data from anaesthetised mice suggest that earlier components are influenced more by adaptation and the physical properties of stimuli, whereas later components are context-dependent (Chen et al., 2015; O’Reilly, 2019a; O’Reilly and Angsuwatanakul, 2021). It has also been shown that this late component in anaesthetised mice relies on intact NMDA receptor function (Chen et al., 2015), comparable with human MMN (Avissar and Javitt, 2018; Rosburg and Kreitschmann-Andermahr, 2016). Curiously, obligatory components of the auditory response generally occur earlier than those recorded in humans, consistent with the theory that sensory response latencies are determined by anatomical size and complexity (Itoh et al., 2022; Javitt et al., 1992; Komatsu et al., 2015), whereas components associated with auditory novelty occur at comparatively similar latencies, apparently contradicting this theory. As such, the relationships between physiology and cortical auditory-evoked potential components in different species remain an unsolved puzzle.

Auditory novelty responses have been studied extensively in basic, clinical, and preclinical animal investigations, although despite this there remains substantial debate concerning their underlying neurophysiology (May, 2021; May and Tiitinen, 2010; Näätänen et al., 2005; O’Reilly et al., 2021). This stems from confounds or alternative explanations for differences between responses to sequences of physically different stimuli that have been challenging to completely dissociate, such as adaptation (May, 2021; May and Tiitinen, 2010) or the inherent modulation of overlapping ERP components by the physical properties of sounds (O’Reilly et al., 2021; O’Reilly and Conway, 2021; Takegata et al., 2008; Todd et al., 2008). In the present study, we revisit data recorded from anaesthetised mice to ask how cortical auditory reflexes in this preparation encode violations of sensory expectations during passive frequency and intensity oddball paradigms. To support this, multivariate pattern analysis (MVPA) and hierarchical recurrent neural network (RNN) modelling are adopted to characterise cortical signal dynamics associated with auditory novelty responses. The MVPA approach was selected to explore subtle effects of stimulus conditions that may be missed by conventional methods (Bae et al., 2020), whereas modelling with a hierarchical RNN provides a tool for studying potential computational principles that underlie the generation of cortical auditory-evoked responses (Barak, 2017; Barrett et al., 2019; Yang and Molano-Mazón, 2021).

## 2. Methods

### 2.1. Data

The data used in this study have been described elsewhere (O’Reilly, 2019a). These experiments were approved by the Animal Welfare and Ethical Review Body, University of Strathclyde, and performed in accordance with the UK Animals (Scientific Procedures) Act 1986.

To summarise, 14 urethane-anaesthetised mice were presented with frequency and intensity oddball sequences while epidural field potentials were recorded bilaterally from electrodes positioned above their primary auditory cortices (as shown in Fig. 1 of O’Reilly, 2019b). In both of the oddball paradigms, standard stimuli were 100 ms, 10 kHz, 80 dB, monophonic pure-tones. In the frequency oddball paradigm, deviant stimuli deviated by ±2.5 kHz, and in the intensity oddball paradigm they deviated by ±10 dB, above and below the standard. These were played with a constant offset to onset inter-stimulus interval of 450 ms. Each sequence included 800 standards (S), 100 increasing deviants (D1) and 100 decreasing deviants (D2). To balance the numbers of trials, standards preceding deviants were extracted, reducing the number of standard trials to 200 for each animal.

**Figure 1.**
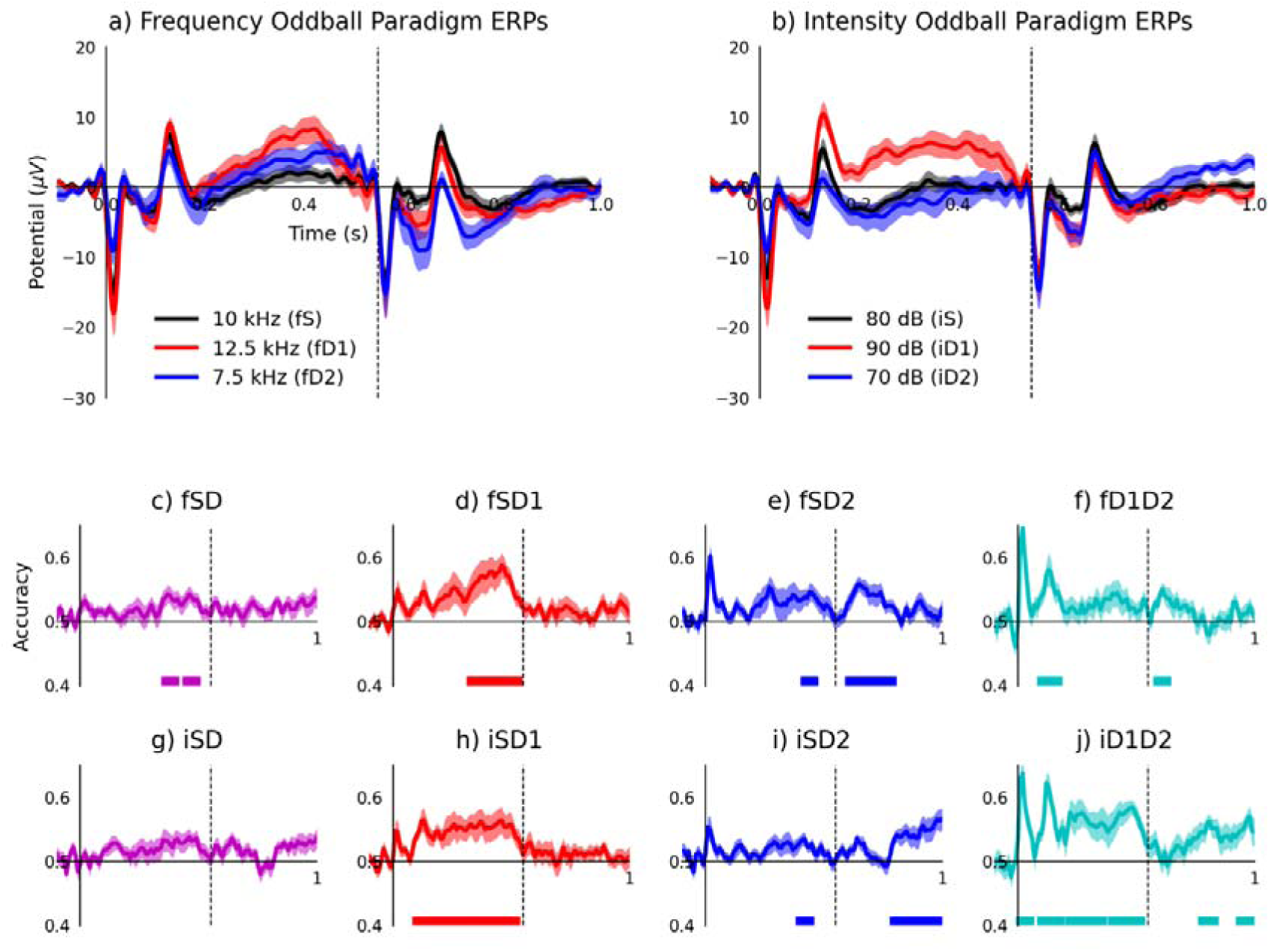
Tone frequency changes and rising sound-level transitions trigger positive long-latency response. a) Grand-average ERP waveforms from frequency oddball paradigm stimuli. b) Grand-average ERP waveforms from intensity oddball paradigm stimuli. c-f) Decoding accuracy for frequency oddball paradigm stimuli: standard vs. deviant (fSD), vs. ascending deviant (fSD1), vs. descending deviant (fSD2), and ascending vs. descending deviant (fD1D2). g-j) Decoding accuracy for intensity oddball paradigm stimuli: standard vs. deviant (iSD), vs. louder deviant (iSD1), vs. quieter deviant (iSD2), and louder vs. quieter deviant (iD1D2). Dashed vertical lines at 0.55 s indicate onset of the second (standard) stimuli. Coloured bars at the bottom of each plot represent statistically significant decoding accuracy (p < α; corrected for multiple comparisons).

Signals were recorded at a frequency of 1000 Hz and subsequently band-pass filtered between 0.1 and 30 Hz. Trials containing two consecutive stimulus responses were extracted; i.e. either responses to two repeated standard stimuli, or responses to an unexpected deviant followed by a standard stimulus, as described previously (O’Reilly, 2019a; O’Reilly and Angsuwatanakul, 2021). These segments spanned from 0.1 s before to 1 s after the first stimulus, capturing two consecutive auditory-evoked responses. Pre-first-stimulus average baseline correction was applied to each trial, and signals were resampled to 100 Hz for MVPA and modelling.

### 2.2. ERP Decoding

#### 2.2.1. Time-wise

To perform MVPA on ERPs, also referred to as ERP decoding (Grootswagers et al., 2017), data from each animal were analysed separately before averaging the results. A linear support vector machine (SVM) was trained to perform binary classifications between ERPs at each time-point, down-sampled to 100 Hz. Four different binary classifications were assessed: standards or deviants (S vs. D; fSD or iSD), standards or increasing-deviants (S vs. D1; fSD1 or iSD1), standards or decreasing-deviants (S vs. D2; fSD2 or iSD2), and increasing-deviants or decreasing-deviants (D1 vs. D2; fD1D2 or iD1D2).

The number of trials presented were equalised (e.g. in the case of S vs. D2, S was under-sampled randomly in order that both conditions had equal trial numbers), and 10-fold cross-validation was performed. This cross-validation was then repeated five times, each time the under-sampling and fold allocation were performed again. The results across all these were then averaged to get an accuracy measure of the decoding at each time point. The MVPA was performed using MVPA-Light (Treder, 2020) and Fieldtrip (Oostenveld et al., 2011). After calculating the time-wise decoding accuracy for each animal, these data were evaluated using t-test statistics, with control for multiple comparisons using cluster-based permutation tests, with the threshold for statistical significance conventionally set to α = 0.05.

#### 2.2.2. Temporal generalisation method

To evaluate potential correlations and interactions between neural responses occurring at different latencies, time-generalised decoding was performed (King and Dehaene, 2014). The same eight binary classifications (fSD, fSD1, fSD2, fD1D2, iSD, iSD1, iSD2, and iD1D2) were performed separately using a linear SVM, although in the temporal generalisation method the model was trained on data from one time point and then tested across all time points. This produced two-dimensional matrices of decoding accuracy by training and test time for each animal. Otherwise, analysis was as per above.

### 2.4. Modelling

#### 2.4.1. Inputs

Sound waveforms comparable with those presented during the in-vivo experiment were produced with a sampling frequency of 100 kHz. These included standard (S), ascending frequency deviant (fD1), descending frequency deviant (fD2), louder intensity deviant (iD1), and quieter intensity deviant (iD2) stimulus conditions. To simulate sound intensity, amplitudes were normalized to 80 dB (i.e. 80 dB sounds had a peak amplitude of 1). These were converted into the time-frequency domain using the short-time Fourier transform (STFT), producing a representation of cortical input from the ascending auditory pathway (Rahman et al., 2020). The STFT was performed on Hann-windowed segments of 200 samples with an overlap of 100 samples. This evaluated frequencies from 0 to 50 kHz, approximating the spectral hearing range of mice, with linear spacing of 0.5 kHz. Magnitudes of the complex STFT output for each trial type were down-sampled to 100 Hz and provided to the model as input features.

Target outputs for the model were obtained by averaging the trials from all of the animals to produce a single ‘idealised experiment’, depicting the most salient electrophysiological features with reduced noise and variability. Left and right auditory cortex channels were also averaged together. This process generated 200 standard, 100 increasing-deviant, and 100 decreasing-deviant trials from frequency and intensity oddball paradigms, equalling 800 trials in total. Model inputs and target outputs were both formatted as sequences of 111 time-samples for training the model in a supervised learning paradigm.

#### 2.4.2. Model architecture and training

A hierarchical RNN was selected to model auditory-evoked potential data. This may be interpreted as a firing rate model that converges towards one of the possible solutions to the inverse source problem. It had 101 input units, each associated with a frequency component of the STFT output. These input units connected to the first of four hidden layers of 64 recurrent units, whose outputs are determined by inputs from the current time-step and feedback from their outputs in the previous time-step. Recurrent connections allow RNNs to learn from sequences of inputs, and are loosely analogous to feedback connections in biological neural networks. The output layer consisted of a single recurrent unit that produced signals in response to simulated auditory inputs.

Input features were fed through the model and its parameters (weights and biases) were optimized to minimise mean-squared-error (MSE) loss between model outputs and target evoked responses. Adaptive moment estimation (Adam) optimization was used with a learning rate of 0.001, beta-1 equal to 0.9 and beta-2 equal to 0.99. Connection weights between layers were initialized from a Glorot uniform distribution (Glorot and Bengio, 2010) and recurrent weights were initialised as an orthogonal matrix from a normal distribution (Saxe et al., 2013). Generalisation of this model architecture was verified by cross-validating over 10-folds, each with 10% of trials of each stimulus type held back for evaluation. This resulted in average terminal MSE for the training set of 114.2 (1.48 SD) and for the validation set of 109.1 (9.71 SD), demonstrating adequate ability of the model architecture to generalise to withheld data. Five identical models were then trained for 500 epochs and evaluated in terms of MSE and Pearson’s correlation coefficient (r^2^) between model outputs in response to each stimulus condition and their associated grand-average ERP waveforms. The best performing model in terms of minimising MSE and maximising r^2^ was subsequently analysed and used in simulated experiments.

#### 2.4.3. Hidden unit categorisation

After establishing the best performing model, its units were categorised based on their time-domain response properties. Units with peak activation latency before stimulus onset, or with activations exceeding three standard deviations above the mean during the pre-stimulus period and having a post-stimulus peak activity less than half of the pre-stimulus peak activity, were categorised as ‘alpha’ units, as these were found to display periodic activity within the alpha frequency range. Units with peak activity during stimulus-on times were categorised as ‘onset’ units, and those with peak activity up to 50 ms after stimulus-off times were categorised as ‘offset’ units. Units with activity peaking between 50 and 150 ms after stimulus-off times were tentatively categorised as ‘safety’ units, while those that peaked between 150 and 450 ms after the first stimulus were categorised as ‘danger’ units. Rationale for this nomenclature is explained in the discussion. Units that consistently had zero activation across all stimulus conditions were classed as ‘zero’ units. Any remaining units that did not fall within these categories were labelled as ‘other’ units, of which there was only one. This time-domain categorisation procedure was applied to all of the hidden units in response to five stimulus conditions (S, fD1, fD2, iD1, iD2) and the most frequent categorisation across stimulus conditions was selected for each unit.

#### 2.4.4. Principal component analysis

Hidden unit activations were transformed into principal component space to examine their principal modes of variance, providing a means of the analysing model, layer, and unit behaviour. Three different aspects of the data were explored using PCA. 1) Model responses to five stimulus conditions, a 5-by-28416 (4 layers, 64 units, 111 time-samples) matrix, were transformed into two principal components that explained 84% of the variance in the data. 2) Layer responses, a 4 × 35520 (5 stimulus conditions, 64 units, 111 time-samples) matrix, were transformed into two components that accounted for 97% of the variance in the data. 3) Units in each layer, four 64-by-555 (5 stimulus conditions, 111 time-samples) matrices, were each transformed into two components that accounted for 68%, 41%, 55%, and 39% of the variance in layer 1, 2, 3, and 4, respectively. Only two principal components were selected for effective visualisation.

#### 2.4.5 Entropy analysis

Sample entropy was calculated from model hidden unit activation signals in response to different stimuli using the method proposed by (Richman and Moorman, 2000). Higher sample entropy is interpreted to reflect greater levels of information production in a dynamic system, while lower sample entropy reflects the opposite. This is another perspective on the conventional view of entropy as being proportional to disorganisation, given that disorganised (changing) signals potentially contain more information than completely organised (unchanging) signals. Entropy measures have been used successfully to describe biological neural signals (Angsuwatanakul et al., 2020; Phukhachee et al., 2019), thus this technique is appropriate for characterising the behaviour of artificial neural signals generated by the model.

#### 2.4.6. Simulated experiments

Audio waveforms of stimuli varying in duration, frequency, and intensity were generated for use in simulated experiments. Different duration stimuli (100, 200, 300, 400, and 500 ms) were all 10 kHz, 80 dB pure-tones; frequency stimuli (5, 7.5, 10, 12.5, and 15 kHz) were 100 ms duration and 80 dB intensity; and intensity stimuli (60, 70, 80, 90, and 100 dB) were 100 ms, 10 kHz pure-tones. Each simulated tone was converted into a time-frequency domain representation, as described above, and applied as model input to examine whether the resulting output would meet logical expectations based on neurophysiological findings reported in the literature.

#### 2.5. Software

Python 3, MNE 0.23.4, Scikit-learn 0.22.2, and Tensorflow 2.4.1 were used for data processing, MVPA and RNN modelling. The Matlab toolboxes Fieldtrip and MVPA-Light were also used to perform statistical analyses.

## 3. Results

### 3.1 Tone frequency changes and rising sound-level transitions trigger positive long-latency response

Decoding of stimulus conditions from ERP waveforms evoked during frequency and intensity oddball paradigms is shown in Figure 1. Grand-average ERP waveforms evoked by frequency oddball paradigm stimuli are plotted in Figure 1a, and those from the intensity oddball sequence are plotted in Figure 1b. The obligatory auditory response is characterised by a negative stimulus-onset peak and a positive stimulus-offset peak, both of which are influenced by the physical properties of eliciting stimuli, and are evoked irrespective of stimulus context (e.g. whether expected or unexpected). Context-dependent, long-latency components of the ERP waveform are also apparent. Both frequency and increasing intensity deviant stimuli induce a positive amplitude feature that peaks at approximately 0.3 to 0.5 s. This is followed by a negative-going feature that coincides with presentation of the following standard stimuli at 0.55 s through to approximately 0.8 s. Responses to standard stimuli that follow quieter deviant stimuli, shown in Figure 1b, also appear to exhibit a positive-going trajectory, resembling the long-latency response evoked by frequency or louder deviant stimuli, which was not recognised in previous analysis of these data. Statistics related to this qualitative description can be found in (O’Reilly, 2019a).

Accuracy of decoding stimulus conditions from responses to frequency oddball paradigm stimuli reinforces observations from grand-average ERP waveform morphologies. Responses to standard stimuli compared with those to both frequency deviants (fSD; Figure 1c) demonstrate significant decoding accuracy over 0.35 to 0.41 s (p = 0.031) and 0.44 to 0.50 s (p = 0.026), aligning with the positive long-latency component. This corresponds to overlapping latencies of significant decoding accuracy in response to ascending (fSD1: 0.32 to 0.54 s, p = 0.001; Figure 1d) and descending frequency deviants (fSD2: 0.41 to 0.47 s, p = 0.037; Figure 1e); although the subsequent negative amplitude feature in descending frequency deviant trials also produced significant decoding accuracy from 0.60 to 0.80 s (p = 0.002). Despite their somewhat similarity trajectories, ascending and descending frequency deviant responses are decoded with significant accuracy (fD1D2; Figure 1f) extending over the stimulus-offset response peak latency, from 0.09 to 0.18 s (p = 0.003) and again from 0.58 to 0.64 s (p = 0.014).

Comparable MVPA applied to stimulus conditions in the intensity oddball sequence reveals a more complex pattern of responses. Starting with the two deviants (iD1D2; Figure 1j), regions of significant decoding accuracy occur during stimulus-onset (0 to 0.06 s, p = 0.023) and stimulus-offset (0.09 to 0.19 s, p = 0.005) response peak latencies, the positive portion of the long-latency response evoked by louder deviants (0.21 to 0.37 s, p = 0.004; 0.39 to 0.53 s; p = 0.001), and the late positive amplitude response evoked by standards following quieter deviants (0.77 to 0.84 s; p = 0.022; 0.93 to 1.0 s, p = 0.025). Relative to the standard, louder deviant stimuli produced significant decoding accuracy between stimulus-offset through to presentation of the second standard stimulus (iSD1: 0.09 to 0.53 s, p < 0.001; Figure 1h). In contrast, quieter deviant stimuli produced a region of significant decoding accuracy near the long latency response to the quieter tone (0.39 to 0.45 s, p = 0.046) and later after the return to the standard tone from 0.79 to 1.0 s (p < 0.001). It is noteworthy that grand-average ERP waveforms (Figure 1b) indicate that this late response is caused by the 80 dB standard tone that follows the 70 dB deviant. These distinctions can help to explain the pattern of significant decoding accuracy between standard and deviant conditions in the intensity oddball paradigm (iSD; Figure 1g), which consists of a relatively short burst from 0.35 to 0.41 s (p = 0.031) and 0.44 to 0.50 s (p = 0.026), influenced predominantly by the louder deviant condition.

### 3.2 Intensity oddball stimuli influence the auditory response asymmetrically

Generalised decoding analysis is presented in Figure 2. Performance of classifying stimulus conditions based on responses evoked by frequency oddball paradigm stimuli (Figure 2a-d) largely reinforce findings from ERP decoding analysis presented in Figure 1c-h, with regions of significant decoding accuracy correlating with long-latency features of the ERP. The seemingly biphasic nature of these long-latency components is highlighted by opposing changes in decoding accuracy when trained and tested over time-points within the ranges of 0.3 to 0.5 s and 0.6 to 0.8 s. For example, when samples from the 0.3 to 0.5 s range are used for training, this increases decoding accuracy above chance for testing over the same interval, while simultaneously driving decoding accuracy below chance over the 0.6 to 0.8 s window, and vice versa. This reflects the positive amplitude shift evoked by frequency-deviant stimuli and the negative amplitude shift that occurs following the subsequent standard stimulus. Generally, significant decoding comparing standard to all deviant stimuli occurred from 0.37 to 0.53 s in training times, and 0.31 to 57 s in the test times (p = 0.024), similar to in the standard versus higher frequency deviance occurring from 0.35 to 0.53 s for the training times and 0.31 to 0.60 s for the test times (p = 0.022). Decoding differentiating the standards to the lower frequency tones was somewhat later, with significant training times occurring from 0.64 to 0.80 s and testing times from 0.60 to 0.82 s (p = 0.012).

**Figure 2.**
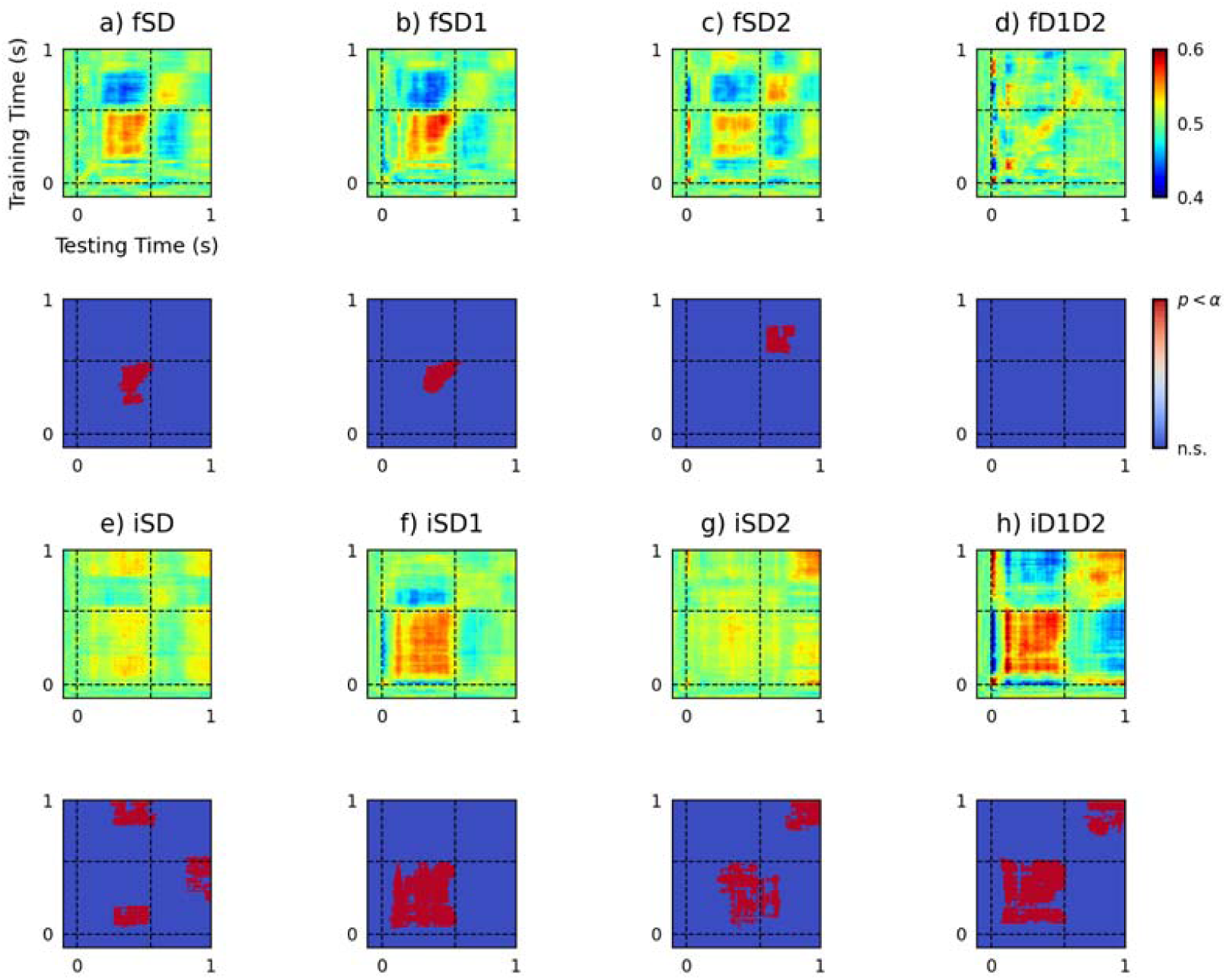
Intensity oddball stimuli influence the auditory response asymmetrically. The top two rows (a-d) display results of pairwise decoding between stimulus conditions in the frequency oddball paradigm. Both frequency deviant stimuli produced significant decoding accuracy during the long-latency window when contrasted with the standard, and were comparatively similar. Decoding accuracy of the descending frequency deviant (fD2) is more pronounced and crosses-over into the second stimulus response. The bottom two rows (e-h) show the results of applying this analysis to intensity oddball paradigm data. The two intensity-deviant stimuli also produced significant decoding accuracies within the window of the first stimulus response, although by inspecting the ERPs (Figure 1b) it can be seen that this was due to amplitude shifts in opposite directions, which nullified decoding accuracy between standards and deviants (iSD) over the first stimulus response window. There is also a late portion of significant decoding accuracy cause by the iD2 condition, which is presumably caused by the sound-level transition between a 70 dB deviant stimulus and an 80 dB standard stimulus. Stimulus onset times at 0 s and 0.55 s are denoted with dashed lines. In statistical plots below each decoding matrix, red represents statistically significant decoding accuracy following an adjustment for multiple comparisons using cluster-based corrections.

Generalised decoding of stimulus condition from responses to intensity oddball paradigm stimuli (Figure 2e-h) is complicated by asymmetries between louder and quieter deviant sounds, and their influence on the sound-level transition between the first and second stimuli in each tone-pair. These complications are emphasised by statistically significant off-diagonal effects when decoding between standard and deviant stimuli (iSD: training = 0.82 to 1.0 s; 0.80 to 0.21 s; 0.25 to 0.59 s, testing = 0.27 to 0.57 s; 0.28 to 0.54 s; 0.82 to 1.0 s; p = 0.033, 0.046, 0.048, respectively; Figure 2e). Note, these off-centre effects are particularly notable as there was no significant decoding in the standard time decoding of these stimuli. Regions of significant decoding accuracy between standard and louder intensity deviants (iSD1; training = 0.40 to 0.54 s, testing = 0.06 to 0.57 s, p = 0.002, Figure 2f), and standard and quieter deviant (iSD2; training = 0.05 to 0.54 s, 0.78 to 1.0 s, testing = 0.23 to 0.71 s, 0.75 to 1.0 s; p = 0.012, 0.038, respectively; Figure 2g) in time-points covering the range of approximately 0.05 s to 0.5 s are thought to result from deviant stimuli evoking amplitude shifts in opposite directions, thereby cancelling each other out when pooled together (Figure 2e). Furthermore, trials where quieter deviants are immediately followed by an increasing sound-level transition (i.e. the following standard, which was 80 dB), produced statistically significant, off-diagonal activity when trained on early time-samples coinciding with the long latency response and tested on onset response activity from the second tone, as seen in Figure 2g. Note this is part of the early cluster in iSD2. There is also significant later decoding in the later time point in the iSD2 condition. This is considered to result from relative differences in amplitude between standard and quieter deviant trials during these distant time-windows that are predictive for stimulus condition. Finally, comparing the two deviant trials (iD1D2) resulted in significant early (training = 0.08 to 0.57 s, testing: 0.06 to 0.57 s, p < 0.001) and later (training = 0.74 to 1.0 s, testing = 0.70 to 1.0 s, p = 0.038) time points.

### 3.3 Hierarchical recurrent neural network fitted to idealised mouse model cortical auditory evoked response

Hierarchical RNNs were trained to generate output sequences matching cortical evoked potentials in response to simulated auditory inputs computed from a STFT. This process is illustrated in Figure 3a. By minimising MSE loss, model outputs became strongly correlated with grand-average ERP waveforms. Learning curves from training five models, shown in Figure 3b, converge towards a common point, comparable with model performance in cross-validation. After training, the best model was selected based on evaluation between its outputs in response to the five stimulus conditions (S, fD1, fD2, iD1, and iD2) and grand-average ERP waveforms. Model 4 outputs had the highest correlation (r^2^ = 0.977, σ = 0.013; Figure 3c) and lowest error (MSE = 0.703, σ = 0.153; Figure 3d) measured across stimulus conditions.

**Figure 3.**
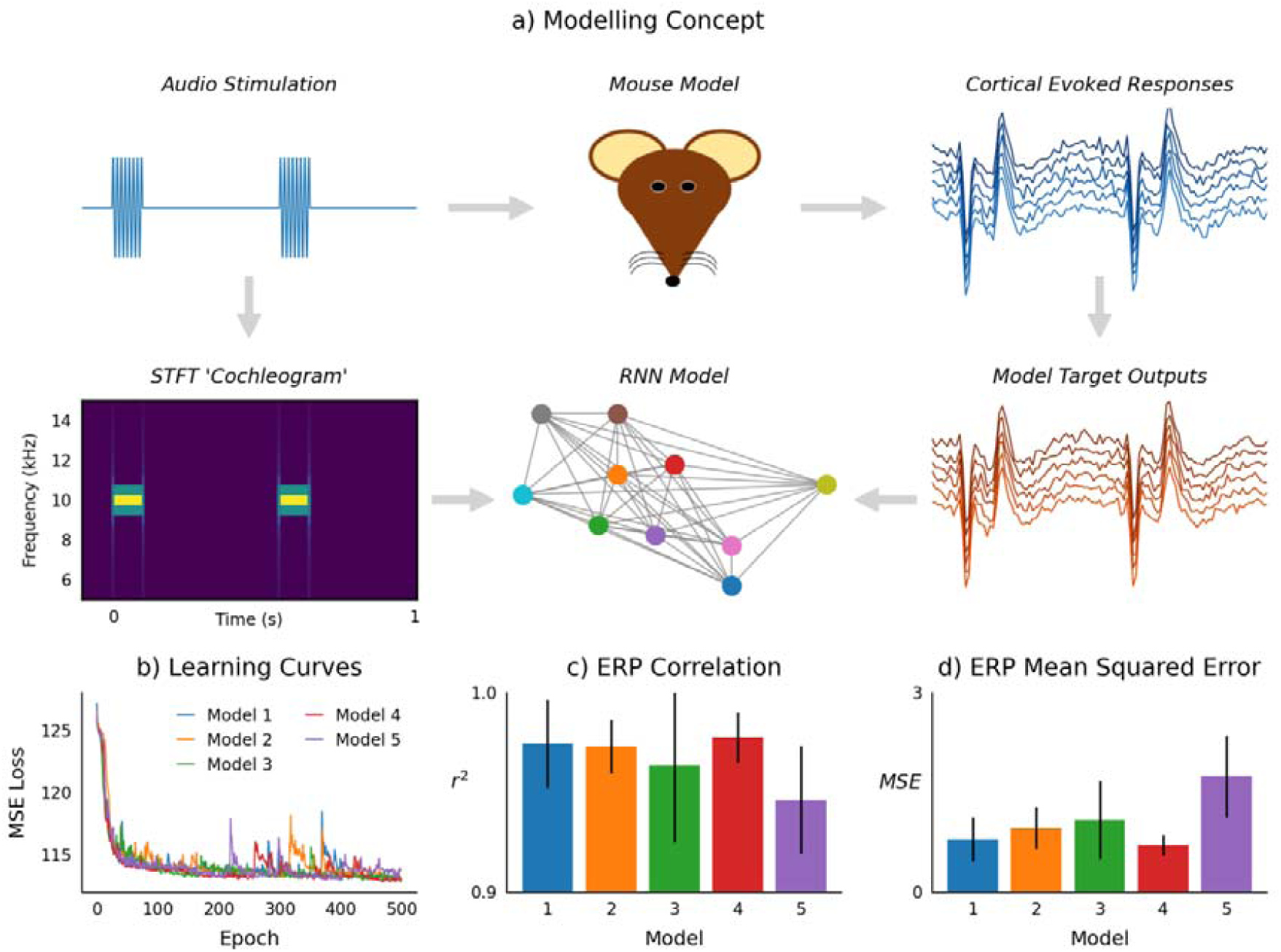
Hierarchical recurrent neural network fitted to idealised mouse model cortical auditory evoked response. a) Data came from an in-vivo experiment in which audio stimulation was applied to the mouse model while recording cortical evoked responses. Sound waveforms were transformed into time-frequency domain ‘cochleograms’ using the STFT, and used to train an RNN model to generate signals equivalent to target outputs from the mouse cortical evoked responses. The RNN model graphic is illustrative, as it actually consisted of four hidden layers, each with 64 units, and a single output unit. b) Learning curves from five models trained over 500 epochs. c) Correlation, and d) MSE, between model outputs and grand-average ERP waveforms from five stimulus conditions (S, fD1, fD2, iD1, iD2); from this analysis, model 4 was identified as the best model. Bar charts display mean with standard deviation.

Best model hidden unit activations and outputs in response to input stimuli are displayed in Figure 4. The upper four rows represent hidden unit activations, while the lower row displays model outputs alongside grand-average ERP waveforms, representing the ground truth. Light colouring in the hidden layer plots reflect higher unit activity. Layers 1 and 4 display several units with notably phasic activations, although in layer 1 these are non-specific, whereas in layer 4 they appear to be context-dependent, occurring during the positive component of the long-latency response evoked by ascending frequency (Figure 4b), descending frequency (Figure 4c), and louder intensity (Figure 4d) stimuli. Across all layers and stimulus conditions there are pronounced unit activations time-locked to auditory input, analogous to in-vivo recordings of biological neurons of the auditory cortex in response to sound stimulation.

**Figure 4.**
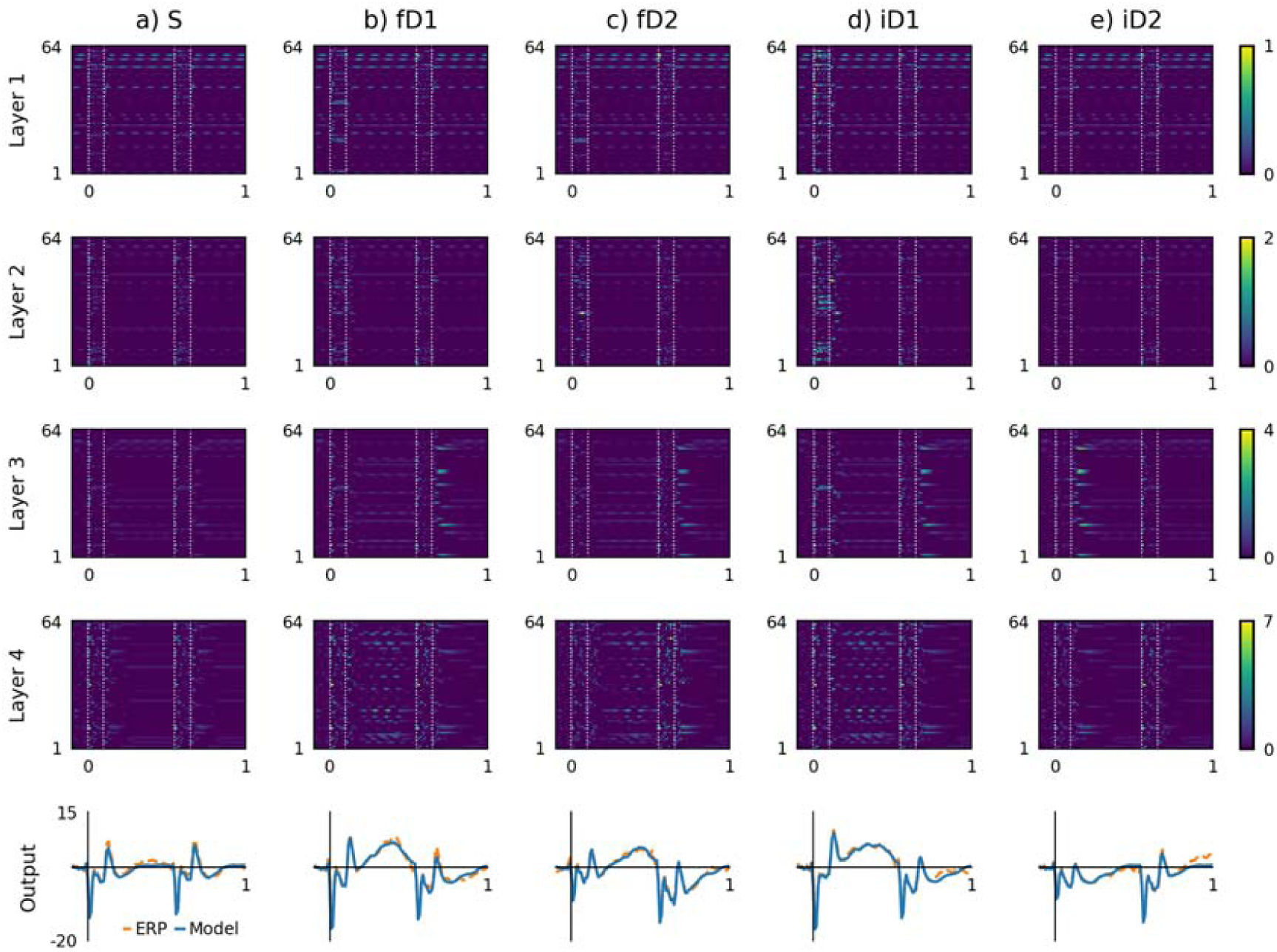
Model hidden unit activations and outputs likened to neural sources. a) Responses to standard stimuli (S). b) Responses to ascending frequency stimuli (fD1). c) Responses to descending frequency stimuli (fD2). d) Responses to higher intensity stimuli (iD1). e) Responses to lower intensity stimuli (iD2). Stimulus onset and offset times are annotated with vertical white dotted lines on the unit activation plots; the second stimulus in each condition was a standard. The model outputs generally match the features of grand-average ERPs; however, they do not capture the positive amplitude response towards the end of the iD2 condition caused by an increasing sound level transition introduced by the second stimulus.

More detail is apparent from the time-domain dynamics of hidden unit activations plotted in Figure 5. Traces are coloured to reflect their categorisation based on temporal response fields, as described in the methods section and outlined in Table 1. This highlights periodic activity of units repeating at approximately 10 Hz, labelled as ‘alpha’ units, as this lies within the alpha range of EEG signal frequencies. Phasic activations are pronounced in layer 1, and are present to a lesser degree in layers 2 and 3. The majority of units are categorised as ‘onset’ units, because their activity peaked during the envelope of auditory stimulation. Together these reliably capture the influence of stimulus frequency and intensity on stimulus onset responses observed in the training data. Comparatively fewer units are categorised as ‘offset’ units, which are responsible for reproducing the offset response feature of the ERP waveform, also present in all of the layers. In contrast, units with peak activity inside the time-window of the positive amplitude long-latency response, categorised as ‘danger’ units, are only present in layer 4. Similarly, although seemingly encoding the opposite phenomena, units with peak activity within the latency of the negative amplitude feature following quieter deviants or transitions from deviants to standards are categorised as ‘safety’ units, which are present in layers 3 and 4.

**Figure 5.**
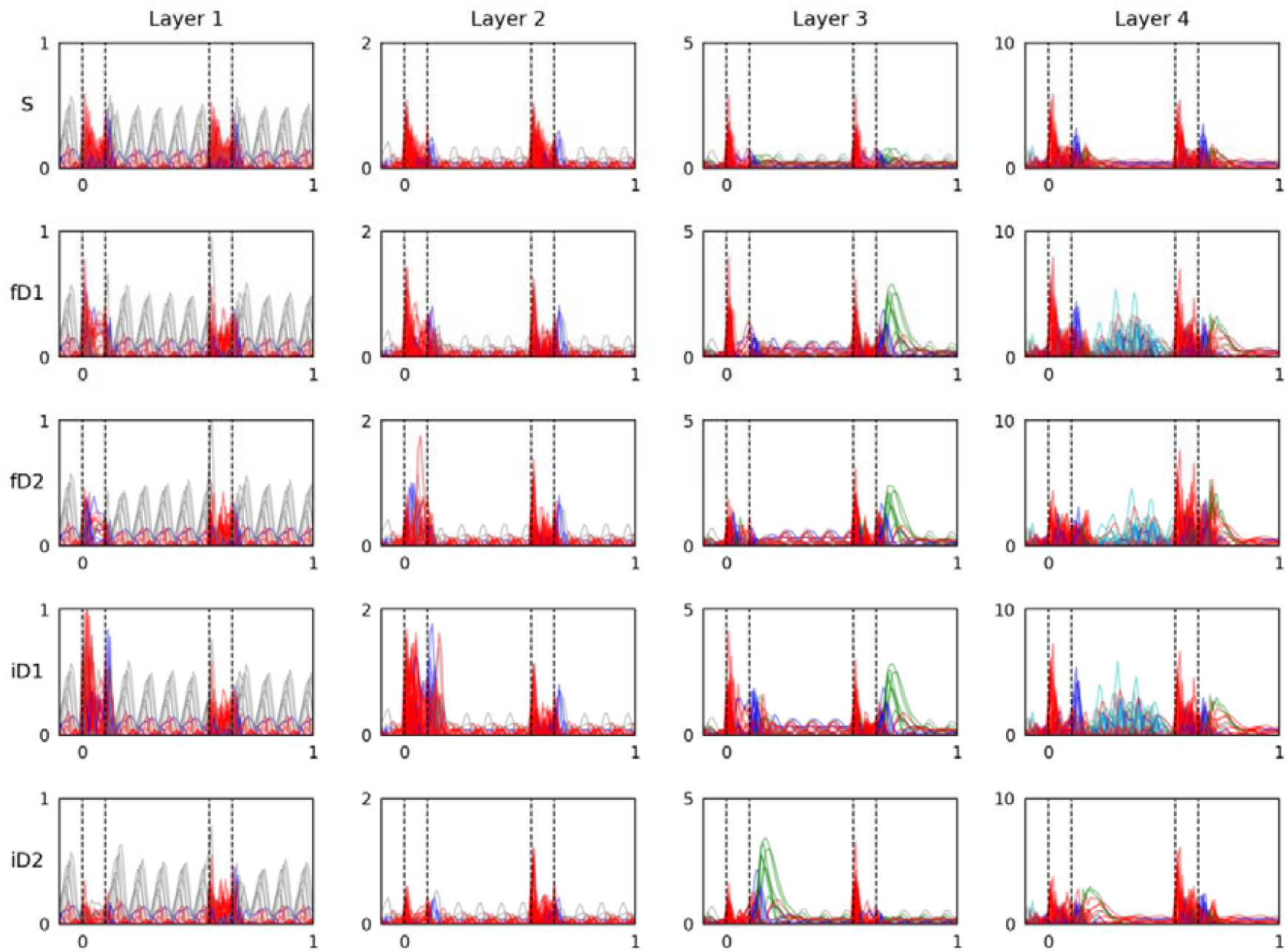
Hidden units classified by their temporal response properties. Five categories of units were defined based on their latency of maximum responsiveness. Units that were highly active during the pre-stimulus baseline period are coloured grey, those whose response peaked during the stimulus-on period are coloured red, those that peaked in the first 50 ms after the stimulus-on period are coloured blue, those that peaked from 50 to 150 ms after the stimulus-on period are coloured green, and those that peaked between 150 and 450 ms after the stimulus-on period are coloured cyan. These five groups are categorised as alpha, onset, offset, safety, and danger, respectively. Remaining units that were unchanging across all stimulus conditions or otherwise did not fall into these categories are coloured black.

**Table 1.** Hidden unit categorisations based on activation peak latency

| Type* | Limits | Layer 1 | Layer 2 | Layer 3 | Layer 4 | Total |
| --- | --- | --- | --- | --- | --- | --- |
| Zero | Unchanging | 11 | 7 | 2 | 3 | 23 |
| Alpha | < 0 s | 11 | 8 | 7 | 5 | 31 |
| Onset | 0-0.1s<br>0.55-0.65 s | 32 | 40 | 30 | 34 | 136 |
| Offset | 0.1-0.15 s<br>0.65-0.7 s | 10 | 9 | 15 | 10 | 44 |
| Safety | 0.15-0.25 s<br>0.7-0.8 s | 0 | 0 | 9 | 4 | 13 |
| Danger | 0.25-0.55 s | 0 | 0 | 0 | 8 | 8 |
| Other | otherwise | 0 | 0 | 1 | 0 | 1 |
| **Total** | - | 64 | 64 | 64 | 64 | 256 |

\*Most frequent categorisation across all stimulus conditions

### 3.4 Principal component analysis groups model stimulus responses, layers, and hidden units by temporal classification

To further examine model behaviour, hidden unit activations were transformed into principal components, presented in Figure 6. Activations of all units in response to the five stimulus conditions represented in principal component space (Figure 6a) exhibit separation between stimuli that evoked a positive amplitude response from 0.3 to 0.5 s (i.e. fD1, fD2 and iD1) and those that did not (i.e. S and iD2). Principal components of activations grouped by layer (Figure 6b) demonstrate the relative similarity between more superficial layers 1 and 2, and dissimilarity between those and deeper layers 3 and 4. Moreover, individual units in each layer transformed into principal components (Figure 6c-f) demonstrate partial clustering corresponding to unit categorisations based on temporal response fields, suggesting that this relatively simple method of categorisation captures the principal modes of variance in unit behaviour.

**Figure 6.**
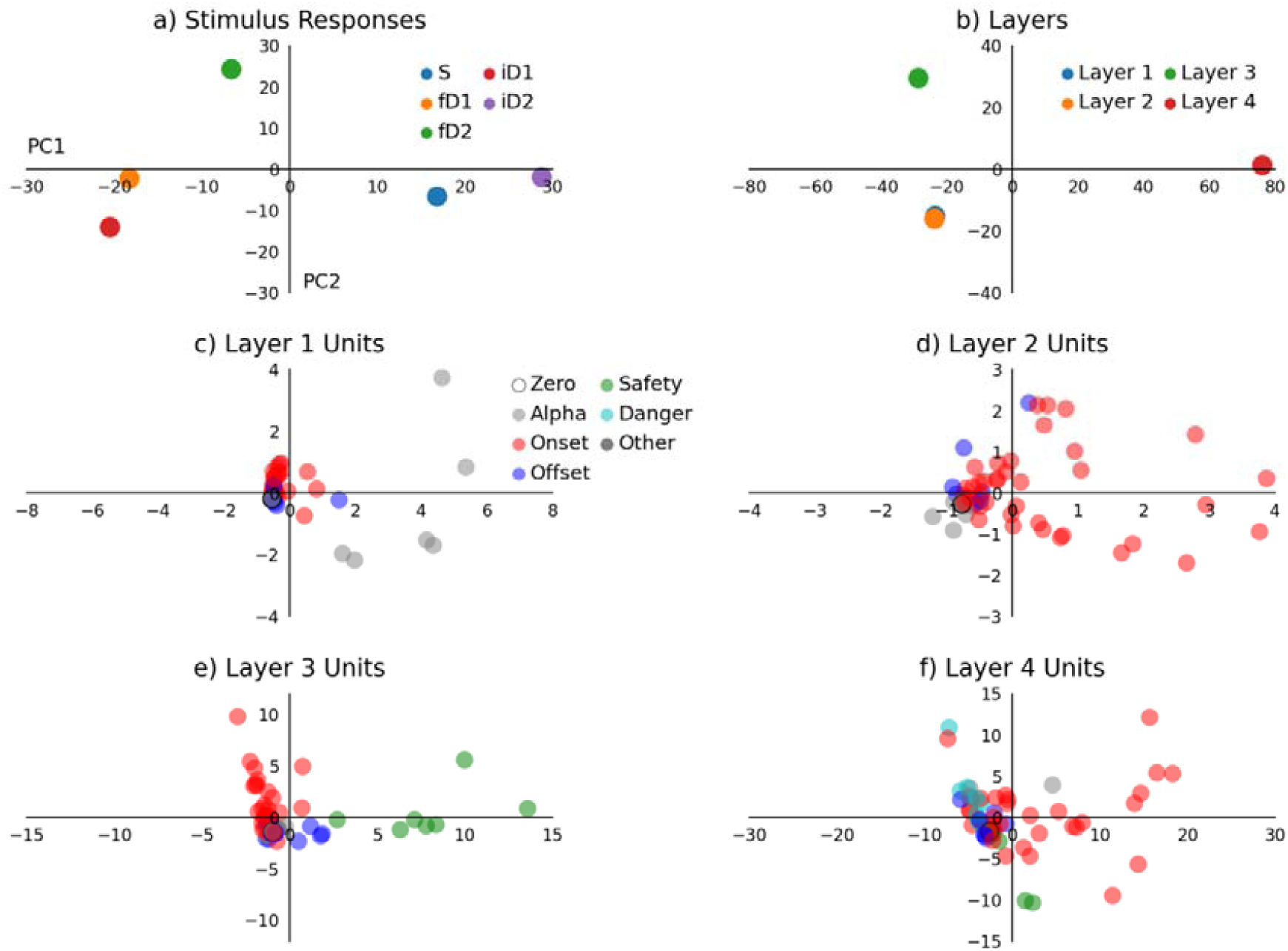
Principal component analysis groups model stimulus responses, layers, and hidden units by temporal response classification. a) Whole-model responses to five stimulus conditions. This shows clear separation between responses to stimuli that did (fD1, fD2, and iD1) and did not (S and iD2) produce a positive amplitude long-latency response over 0.3 to 0.5 s post-stimulus. b) Responses of model layers across all stimulus conditions. Layers 1 and 2 appear to be very similar, whereas layers 3 and 4 account for most of the variance in the first two principal components. Responses of layer 1-4 hidden units are plotted in (c-f), respectively. In (c-f), units are coloured according to their categorisation based on activation peak latency. Time-domain categorisations reflect partial clustering of units in two-dimensional principal component space.

### 3.5 Interaction between superficial and deep layer activations with environmental salience of stimulus conditions

Hidden units that did not exhibit changes in activity had zero entropy, thus containing no information. Across model layers, the numbers of units that had zero entropy in response to different stimulus conditions are plotted in Figure 7a. The first and second layers had more units with zero entropy. From the units with nonzero entropy, layer average entropy during five input stimulus conditions are given in Figure 7b. This shows relatively high entropy in active units of the first layer across all stimulus conditions. There appears to be an interaction between having fewer zero-entropy units in layer 2 and higher mean entropy in layer 4 during frequency and increasing intensity deviant stimuli. Median sample entropy of units in layers 1 to 4 were 0.060, 0.070, 0.101, and 0.164, respectively; although this was clearly influenced by the numbers of zero-entropy units in each layer. Median sample entropy of stimulus conditions S, fD1, fD2, iD1, and iD2 were 0.090, 0.107, 0.110, 0.107, and 0.084, respectively. Average sample entropy of categorised units was also calculated as follows: zero units (n = 23, mean = 0.0, σ = 0.0), alpha units (n = 31, mean = 0.251, σ = 0.283), onset units (n = 136, mean = 0.192, σ = 0.157), offset units (n = 44, mean = 0.139, σ = 0.134), safety units (n = 13, mean = 0.123, σ = 0.103), danger units (n = 8, mean = 0.165, σ = 0.058), and other units (n = 1, mean = 0.052, σ = 0.0).

**Figure 7.**
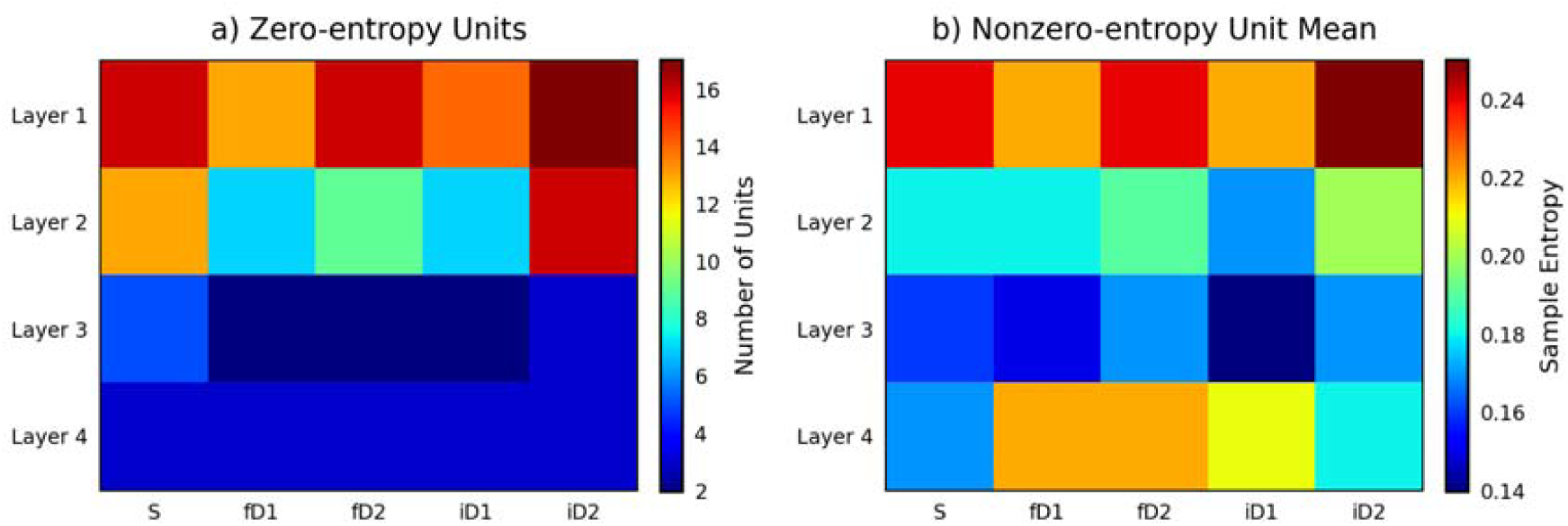
Interaction between superficial and deep layer activations with environmental salience of stimulus conditions. a) The numbers of unresponsive units that showed zero entropy across layers and stimulus conditions. This data presents a layer-wise descending gradient, with fewer unresponsive units in deeper layers. There also appears to be a marginal stimulus-condition effect, with fD1, fD2, and iD1 inputs causing relatively fewer zero-entropy units in layers 2 and 3 than S or iD2 stimuli. b) Average sample entropy from the remaining nonzero-entropy units in each layer in response to different stimulus conditions. Layer 1 units exhibit higher average sample entropy, with a tendency towards decreasing entropy in deeper layers, with the exception of a sharp increase in response to stimuli that evoked a positive amplitude long-latency response (i.e. fD1, fD2 and iD1). This analysis tentatively hints towards a possible link between the activity of fewer nonzero-entropy units in layers 2/3 and downstream initiation of context-dependent signals observed in the activity of layer 4 units.

### 3.6 Simulated auditory inputs elicit stereotypical responses to tone duration and intensity, but not frequency

Simulated auditory inputs not included among the set of training stimulus conditions were applied to the model to simulate experiments, the results of which are displayed in Figure 8. These were evaluated qualitatively based on knowledge from prior neurophysiological investigations (e.g. O’Reilly and Conway, 2021). Different duration stimuli produced stimulus offset responses with peak latency positively correlated with stimulus duration, in agreement with expectations. Different frequency tones influenced stimulus onset and offset response peak amplitudes, although not in the proportional manner expected based on prior results. Moreover, only deviant frequencies presented during model training elicited long-latency responses, defying the logical assumption that other frequencies which differ from the standard would also produce similar long-latency responses in-vivo. Different intensity stimuli influenced stimulus onset and offset response peak amplitudes in a proportional manner, and both louder stimuli generated positive amplitude long-latency responses, meeting expectations.

**Figure 8.**
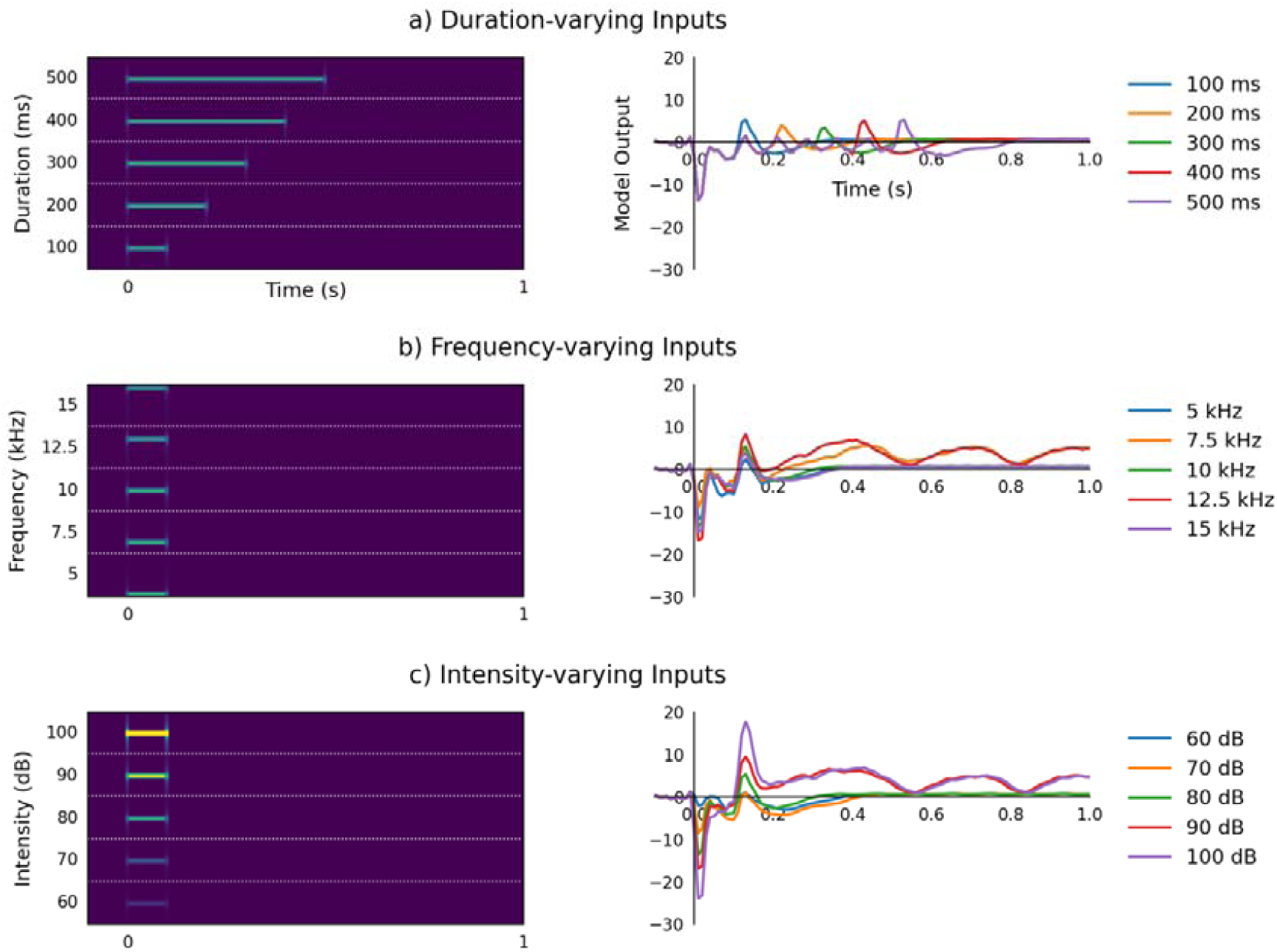
Simulated auditory inputs elicit stereotypical responses to tone duration and intensity, but not frequency. a) Five different duration sounds were simulated and applied as input to the trained RNN (left side), producing simulated ERP waveforms (right side). These waveforms are comparable with those observed in (O’Reilly and Conway, 2021).b) Responses to five different frequency sounds were examined. Here the expected relationship between stimulus onset response peak amplitude and tone frequency is not observed, and a long-latency response is only evoked by frequencies of deviant stimuli used for training the model. c) Responses to five different intensity sounds were also examined. These exhibit correlation between sound intensity and stimulus onset and offset response peak amplitudes, as expected. Moreover, both of the louder tones evoked positive amplitude long-latency responses, whereas the quieter tones evoked more subtle negative amplitudes comparable with those of the standard stimulus. In left-hand side panels, horizontal white dotted lines separate different input stimuli, which are shown from 4.5 to 15.5 kHz.

## 4. Discussion

### 4.1 Tone frequency changes and rising sound-level transitions trigger positive long-latency response

This discussion with concentrate on novel findings revealed from MVPA and analysis of the fitted hierarchical RNN model. However, to briefly summarise, previous analysis of these data using a double-epoch subtraction found that ascending and descending frequency, and louder but not quieter, novel sounds evoke a long-latency mismatch response from anaesthetised mice (O’Reilly, 2019a). As such, this long-latency mismatch response may be thought to reflect environmental salience, not an indiscriminate change-detection mechanism. Something overlooked previously, but brought to the fore by the results of MVPA in Figure 1 and Figure 2, is that standard stimuli following quieter deviant stimuli (i.e. 70 dB to 80 dB) trigger a positive amplitude long-latency response resembling that evoked by increasing intensity deviant stimuli (i.e. 80 dB to 90 dB). Conversely, decreasing sound-level transitions (i.e. either 80 dB to 70 dB or 90 dB to 80 dB) appear to elicit a more subtle, slightly earlier, negative amplitude shift. These observations are consistent with findings from human EEG that demonstrate a prominent role of sound intensity level transition in modulating ERP component amplitudes during the time-window of MMN and P3a (Barry et al., 2022; O’Reilly, 2021a). This adds to evidence suggesting that neural responses to oddball stimulation depend on interactions between physiological adaptation and the physical properties of stimuli, which confound any potential role of prediction-error signalling (Fishman and Steinschneider, 2012; Lazar and Metherate, 2003; May, 2021; May and Tiitinen, 2010; O’Reilly, 2021b; O’Reilly et al., 2021; O’Reilly and Conway, 2021; Solomon et al., 2021; Umbricht et al., 2005).

### 4.2 Negative, ‘safety’, or positive, ‘danger’, long-latency responses can be interpreted to reflect environmentally inconspicuous or salient auditory stimuli

Informed by the results of MVPA and the RNN model, these long-latency responses are referred to using the term ‘danger’ for those that were initiated by an increase in environmental salience of auditory input, such as frequency deviant or increasing intensity deviant stimuli; contrasted with ‘safety’ responses, which were observed following relatively inconspicuous stimuli, such as quieter deviant stimuli or standard stimuli following a danger response. These danger and safety responses are suggested to be distinct from consecutive presentation of two standard stimuli, which reflect neutral environmental salience, and thereby evoke neither safety nor danger responses (top row of Figure 5). While these responses occur during the latency range of human MMN and P3a, it cannot be said definitively that they reflect the same underlying neurophysiological processes, given that the relationships among ERP components, neuroanatomy and complexity of perceptual processing are incompletely understood (Itoh et al., 2022; Javitt et al., 1992; Komatsu et al., 2015). Nevertheless, the findings here replicate those found elsewhere in the animal model literature. Comparable long-latency responses to frequency oddball stimuli have been observed in anaesthetised rodents (Casado-Román et al., 2020; Chen et al., 2015; O’Reilly and Angsuwatanakul, 2021; Ruusuvirta et al., 1998). Curiously though, these have been absent from studies employing similar or near-identical stimulation and recording protocols in conscious mice (Harms et al., 2014; O’Reilly and Conway, 2021). To reconcile this apparent dichotomy between the presence of long-latency mismatch responses in anaesthetised rodents and absence thereof in conscious rodents, it may be speculated that the long-latency response to environmentally salient auditory stimuli quickly habituates if not reinforced in conscious animals, typical of the central orienting reflex associated with noradrenergic neurotransmission mediated via the locus coeruleus (Sara and Bouret, 2012; Shine, 2019). Future studies in anaesthetised and conscious animals may be designed to explore this hypothesis.

### 4.3 Hidden units classified by their temporal response properties

Model hidden units were classified based on temporal response fields, essentially based on their latency of peak activation, into alpha, onset, offset, safety, and danger units. Unresponsive units and units that did not fit into any of these categories were also considered. Time-domain patterns of unit activity in response to different stimulus conditions are shown in Figure 5, with traces coloured according to unit categorisation. The PCA results in Figure 6c-f demonstrate that these classifications reasonably-well capture the primary modes of variance in unit behaviour across all layers. It is evident that the patterns of activity from some of the hidden units spontaneously adopt an alpha frequency oscillation, presenting an emergent parallel with the prominent alpha rhythm observed in human EEG recordings (Quigley, 2021). The activity of these alpha units was suppressed during stimulus presentation, which tangentially lends support for the longstanding phase-reset theory of evoked potentials (Hanslmayr et al., 2007).

### 4.4 Interaction between superficial and deep layer activations with environmental salience of stimulus conditions

Analysis of sample entropy calculated from hidden unit activations shown in Figure 7 hints towards a potential relationship between the behaviour of units in superficial layers with downstream manifestation of long-latency responses in deeper layers. Specifically, there were fewer layer 2 zero-entropy units and higher average sample entropy among layer 4 units during fD1, fD2, and iD1 stimulus conditions, all of which elicited a positive amplitude long-latency response, tentatively suggesting that the trigger signal for this long-latency response originates in earlier layers and feeds through the hierarchical model to influence its output. This offers a small glimpse into the potential structure underlying network responses to contextually different stimuli, however, development of more advanced analytical techniques will be required to explore these relationships more deeply (Barrett et al., 2019; Yang and Molano-Mazón, 2021).

### 4.5 Emergent computational neurophysiology observed from the model

The majority of hidden units were maximally active during the time-window of simulated auditory tones. This parallels biological neurons of the auditory cortex that are excited by sound stimulation (Bajo and King, 2012; King et al., 2018). Second in number were offset response units, for which the existence of biological counterparts is also supported by an extensive literature (Kopp-Scheinpflug et al., 2018; O’Reilly, 2019a; Solyga and Barkat, 2021). Relatively fewer units were grouped into more abstract categories of ‘safety’ and ‘danger’ units; safety units were located in layers 3 and 4, whereas danger units were situated only in layer 4. This somewhat anthropomorphic terminology has been selected to describe the relative environmental salience of stimuli that produced these responses. Danger units were activated by presentation of what are assumed to be salient changes in auditory input, specifically the first stimuli in fD1, fD2 and iD1 conditions. In contrast, comparatively inconspicuous changes in auditory input, such as the first stimulus in the iD2 condition, or second stimuli in fD1, fD2, or iD1 conditions, elicited activity from safety units. The existence of cortical neurons serving comparable functions is less firmly established than the aforementioned parallels between model behaviour and neurophysiological findings, although this seems a reasonable hypothesis derived from the model; particularly given there is evidence of neurons in subcortical structures that encode relative safety or danger attributed to incoming stimuli (Rogan et al., 2005; Sangha et al., 2013). Interestingly, these safety and danger units may also bear some of the hallmarks of negative and positive prediction error neurons that encode opposite direction mismatches between input stimuli and expectations (Hertäg and Clopath, 2022).

An interesting wrinkle exposed by this aspect of model behaviour is that ERP component amplitudes accompanying the activity of safety and danger units are of opposite polarity. Were the biological existence of these units to be confirmed, this would challenge present formulations of the predictive coding theory of auditory processing by implying that prediction errors emerge differently based on stimulus context, at least with respect to environmental salience. Inclusion of this seemingly natural facet of sensory processing should not be at odds with a complete theory of auditory perception. Although, at a minimum this would require revision of the dominant view of predictive coding, which currently lacks an explanation for these observations (Casado-Román et al., 2020; Friston, 2005; Garrido et al., 2009; Lieder et al., 2013). There are alternative, albeit perhaps less exotic or conceptually appealing, explanations that have been proposed to account for differential responses to passive auditory oddball stimulation (Butler, 1968; May, 2021; May and Tiitinen, 2010; O’Reilly, 2021b, 2021a; O’Reilly et al., 2021; O’Reilly and Conway, 2021). However, these are rarely considered preferable to the well-supported predictive coding framework, even when dutifully considered (Lacroix et al., 2022). As they currently stand, neither predictive coding nor alternative sensory processing theories succinctly encapsulate all of the physiological phenomena observed in response to sequences of sounds, which shall therefore remain the purview of future efforts directed towards improving our understanding these processes.

### 4.6 Simulated auditory inputs elicit stereotypical responses to tone duration and intensity, but not frequency

Simulated experiments were conducted as a means of exploring the validity of model behaviour under conditions that were not exposed during training (Wacongne et al., 2012). Figure 8 displays the findings from three such experiments with the best-fitting hierarchical RNN. In these experiments, inputs representing five physically different audio stimuli were constructed and passed through the model to elicit output waveforms that were qualitatively evaluated against expectations based on established neurophysiology. Simulated duration-varying stimuli (Figure 8a) produced close agreement with results from a many-standards control sequence presented to the same cohort of anaesthetised mice (O’Reilly and Conway, 2021), presenting onset and offset responses also seen in anaesthetised rats (Nakamura et al., 2011). This suggests that the trained model reliably captures stimulus onset and offset responses, without producing unexpected activity, in response to simulated audio stimuli with different durations.

Results from simulated frequency-varying inputs (Figure 8b) are further from neurophysiological expectations. The relationships between tone frequency and obligatory stimulus-on and stimulus-off response peak amplitudes were not proportional, therefore not in agreement with previous findings (Nakamura et al., 2011; O’Reilly and Conway, 2021). Furthermore, long-latency responses were only observed from deviant frequency stimuli exposed to during training, defying the expectation that other frequencies that deviate from the standard would also produce comparable responses in-vivo (Casado-Román et al., 2020; Chen et al., 2015; O’Reilly and Angsuwatanakul, 2021). This suggests that the model has not learned to trigger this response to frequencies that differ from the standard, rather it has learned to produce long-latency responses only to the specific deviant frequency stimuli used during training (7.5 and 12.5 kHz). Perhaps training the model across multiple datasets with different stimulus frequencies as standards and deviants would assist in overcoming this limitation (Yang and Molano-Mazón, 2021). Alternatively, training the model on longer sequences spanning several stimuli may be necessary to truly learn the underlying relationship between responses to frequent standard stimuli and infrequent deviant stimuli.

Model outputs in response to different intensity stimuli (Figure 8c) provide closer agreement with physiological expectations. These depict a proportional relationship between simulated stimulus intensity level and the magnitudes of onset and offset responses, which may be expected in-vivo (Bajo and King, 2012; O’Reilly and Conway, 2021). Also, both louder stimuli produced positive amplitude long-latency responses, which appeals to the assumption that environmentally salient stimuli ought to generate similar responses. Quieter stimuli produced a more subtle, negative dip in output waveforms, corresponding to the safety signal already mentioned.

### 4.7 Further Considerations

We ought to acknowledge that any links between model behaviour and neurobiology are necessarily speculative. The present model portrays interesting parallels with experimental neurophysiology in terms of its spontaneous alpha-frequency rhythms, onset and offset units, and potentially also, although yet to be confirmed, danger and safety units, which encode differential responses to environmentally salient and inconspicuous stimuli, respectively. Nevertheless, computational models are treated as tools for studying the potential mechanisms employed within the brain, and are not assumed to be directly homologous. Specifically in relation to artificial neural networks, these are known to be biologically implausible, but can still be valuable for neuroscience research (Barak, 2017; Barrett et al., 2019; Yang and Molano-Mazón, 2021).

One of the particularly appealing elements of artificial neural networks, despite their biological implausibility, is that they develop latent states that may be considered roughly analogous to the behaviour of neural sources, thereby enabling a richer analysis of the potential dynamics of neural activity underlying patterns observed in electrophysiology data. However, it is important to avoid over-interpretation. For example, numbers used to refer to model layers do not imply any hypothetical relationship with cortical layers that are conventionally numbered I to VI. Moreover, the terms superficial and deep, used in the context of artificial neural networks, refer to the relative proximity of layers to the input signal, not to be confused with how these terms are used to discuss cortical layers. It may be possible to apply a vast array of established and novel analytical methods to study the hierarchical RNN, which will provide plenty avenues for future research, although the present article’s scope is limited to keep it concise.

As a final cautionary note, the present application of an RNN to model ‘idealized experiment’ data is subject to some of the same limitations as conventional ERP analysis, such as assuming that evoked components are relatively stationary across subjects and trials. In many cases this may be an invalid assumption. However, MVPA is partially immune to this limitation, being applied to individual subject data separately, and its application has confirmed (as shown in Figure 1 and Figure 2) that the long-latency components evoked by frequency and rising intensity level sounds are not spurious phenomena.

## 5. Conclusion

This reanalysis of cortical auditory-evoked potentials from anaesthetised mice in response to frequency and intensity oddball paradigms offers two related insights that were undiscovered in previous examinations of these data. Firstly, asymmetric sound level transitions in the intensity oddball paradigm modulate cortical electrophysiology in different respects, such that rising sound level transitions, which may be perceived as reflecting greater environmental salience, akin to frequency deviant stimuli, produce a positive amplitude long-latency feature; whereas relatively inconspicuous, falling sound level transitions produce a subtle negative amplitude long-latency feature in the evoked response waveform. Secondly, through the mechanisms depicted by a computational model, these features correlating with relative changes in environmental salience can be conceptualised as reflecting potential ‘danger’ or ‘safety’ responses, respectively, which are encoded by the activity of distinct sources that influence the cortical auditory-evoked response. The hypothesised biological existence of these separate sources may be evaluated in future animal or human neuroimaging experiments. This study also highlights the potential synergy between MVPA and computational modelling approaches, and the myriad intriguing parallels between established auditory neurophysiology and the emergent behaviour of a hierarchical recurrent neural network fitted to cortical auditory evoked potential data.

## Acknowledgements

Data used in this study were obtained during a doctoral training award funded by the United Kingdom Engineering and Physical Sciences Research Council [grant no. EP/F50036X/1]. The computational model was trained using a computer system and graphics processing unit (NVIDIA Titan RTX) purchased with financial support from the Research Institute of Rangsit University [grant no. 90/2561].

## Author Contributions (CRediT Statement)

Jamie A. O’Reilly: Conceptualization, Methodology, Software, Formal analysis, Data Curation, Writing - Original Draft, Writing - Review & Editing, Visualization. Thanate Angsuwatanakul: Formal analysis, Writing - Review & Editing. Jordan Wehrman: Conceptualization, Methodology, Software, Formal analysis, Writing - Review & Editing.

## Data availability

Data and code associated with this article can be accessed from [*will be shared on a public repository prior to publication*].

## Abbreviations

σ: standard deviation
Adam: adaptive moment estimation
D: deviant stimuli
D1: increasing deviant stimulus
D2: decreasing deviant stimulus
EEG: electroencephalography
ERP: event-related potential
fD1: ascending frequency deviant stimulus condition
fD1D2: frequency oddball paradigm D1 vs. D2 decoding
fD2: descending frequency deviant stimulus condition
fSD: frequency oddball paradigm S vs. D decoding
fSD1: frequency oddball paradigm S vs. D1 decoding
fSD2: frequency oddball paradigm S vs. D2 decoding
iD1: rising intensity deviant stimulus condition
iD1D2: intensity oddball paradigm D1 vs. D2 decoding
iD2: falling intensity deviant stimulus condition
iSD: intensity oddball paradigm S vs. D decoding
iSD1: intensity oddball paradigm S vs. D1 decoding
iSD2: intensity oddball paradigm S vs. D2 decoding
MMN: mismatch negativity
MSE: mean squared error
MVPA: multivariate pattern analysis
NMDA: n-methyl-d-aspartate
P3a: involuntary P300 component
PCA: principal component analysis
RNN: recurrent neural network
RON: reorienting negativity
S: standard
STFT: short time Fourier transform
SVM: support vector machine

